# Second Boost of Omicron SARS-CoV-2 S1 Subunit Vaccine Induced Broad Humoral Immune Responses in Elderly Mice

**DOI:** 10.1101/2024.02.05.578925

**Authors:** Eun Kim, Muhammad S. Khan, Alessandro Ferrari, Shaohua Huang, Thomas W. Kenniston, Irene Cassaniti, Fausto Baldanti, Andrea Gambotto

## Abstract

Currently approved COVID-19 vaccines prevent symptomatic infection, hospitalization, and death from the disease. However, repeated homologous boosters, while considered a solution for severe forms of the disease caused by new SARS-CoV-2 variants in elderly individuals and immunocompromised patients, cannot provide complete protection against breakthrough infections. This highlights the need for alternative platforms for booster vaccines. In our previous study, we assessed the boost effect of the SARS-CoV-2 Beta S1 recombinant protein subunit vaccine (rS1Beta) in aged mice primed with an adenovirus-based vaccine expressing SARS-CoV-2-S1 (Ad5.S1) via subcutaneous injection or intranasal delivery, which induced robust humoral immune responses (1). In this follow-up study, we demonstrated that a second booster dose of a non-adjuvanted recombinant Omicron (BA.1) S1 subunit vaccine with Toll-like receptor 4 (TLR4) agonist RS09 (rS1RS09OM) was effective in stimulating strong S1-specific immune responses and inducing significantly high neutralizing antibodies against the Wuhan, Delta, and Omicron variants in 100-week-old mice. Importantly, the second booster dose elicits cross-reactive antibody responses, resulting in ACE2 binding inhibition against the spike protein of SARS-CoV-2 variants, including Omicron (BA.1) and its subvariants. Interestingly, the levels of IgG and neutralizing antibodies correlated with the level of ACE2 inhibition in the booster serum samples, although Omicron S1-specific IgG level showed a weaker correlation compared to Wuhan S1-specific IgG level. Furthermore, we compared the immunogenic properties of the rS1 subunit vaccine in young, middle-aged, and elderly mice, resulting in reduced immunogenicity with age, especially an impaired Th1-biased immune response in aged mice. Our findings demonstrate that the new variant of concern (VOC) rS1 subunit vaccine as a second booster has the potential to offer cross-neutralization against a broad range of variants and to improve vaccine effectiveness against newly emerging breakthrough SARS-CoV-2 variants in elderly individuals who were previously primed with the authorized vaccines.

## Introduction

Vaccination has been a valuable public health strategy for controlling COVID-19 and has greatly reduced the rate of hospitalization, severe disease, and death (2). However, vaccination becomes less effective with increased age, as older individuals exhibit lower serum neutralization and immunoglobulin (Ig)G/A titers after a single vaccination with Pfizer’s BNT162b2 messenger RNA vaccine (3). Furthermore, older individuals or immunocompromised patients have responses that wane more quickly, meaning an increased risk of reinfection over time (4, 5). Moreover, new variants of SARS-CoV-2 continue to emerge and circulate, evading the immune response and leading to alterations in the neutralizing antibody targets, rendering existing vaccines less effective and increasing the risk of breakthrough infections. Notably, the Omicron variant spread globally faster than any previous variant of the SARS-CoV-2 coronavirus, infecting even those who had been vaccinated, regardless of the vaccine type and platform (6–8) or prior COVID-19 infection (9). Indeed, the majority (89.2%) of fully vaccinated patients hospitalized due to the SARS-CoV-2 Omicron variant were over 65 years old and/or severely immunosuppressed, despite the Omicron variant being less virulent than previous strains (10). In-hospital mortality did not significantly differ among patients requiring intensive care, regardless of their vaccine status, with only age showing a significant relationship with higher in-hospital mortality (11). Israeli trial shows a fourth vaccination of ancestral strain raises antibody levels but provides little extra protection against SARS-CoV-2 Omicron infection (12). Therefore, the administration of additional heterologous boosters is recommended to enhance vaccine effectiveness and protect elderly people (13). It is crucial to develop strategies to improve vaccine effectiveness with alternative antigen sequences. Furthermore, in humans, non-inflammatory IgG4 levels increased several months after the second and third mRNA vaccination (14). This class switch was associated with a reduced capacity of the spike-specific antibodies to mediate antibody-dependent cellular phagocytosis (ADCP) and antibody-dependent complement deposition (ADCD), which are critical for antiviral immunity (15). This has led to the consideration of alternative vaccine platforms for boosting.

Nearly all COVID-19 vaccine types target a spike protein, particularly protein subunit vaccines containing specific products of the virus rather than complete viral particles, such as the spike protein or its segments as antigens intended to elicit humoral and cellular immunity and provide effective protection (16). Subunit vaccines of COVID-19 offer several advantages, including suitability for people with compromised immune systems, safety due to the absence of live components, well-established technology, and relative stability (17). Heterologous subunit-based adjuvanted vaccines are known to induce robust and durable immune responses. Many recent studies have shown that spike subunit booster vaccines (e.g., CoV2 preS dTM-AS03, NVX-COV2373/NVX-COV2515, or MVC-COV1901) elicited robust neutralizing antibodies against multiple variants in populations primed with adenovirus-based vaccines (18–22). Moreover, a protein subunit vaccine as a booster showed the best safety profiles in terms of both local and systemic adverse events compared to AZD1222 and mRNA-1273 (19), and three doses of NVX-CoV2373 elicited high neutralizing titers against Omicron BA.1 and BA.4/BA.5, with responses similar in magnitude to those triggered by three doses of an mRNA vaccine (23). Furthermore, the spike protein subunit vaccine demonstrated superior viral control in the upper airway against BA.5 infection compared to the mRNA-1273 vaccine in non-human primates primed with two doses of mRNA-1273, although comparable neutralizing antibodies were induced by both booster vaccines (24).

Previous study from our lab demonstrated that a single immunization with an adenovirus-based vaccine expressing SARS-CoV-2-S1 (Ad5.S1), administered via either subcutaneous (SC) injection or intranasal (IN) delivery, and a recombinant protein subunit vaccine (rS1Beta) booster effectively induced long-lasting and cross-neutralizing antibodies against SARS-CoV-2 variants in aged mice. This indicates that subunit vaccines are considered safe and effective for use as booster vaccines (25). Herein, as ongoing research, we evaluated that the vaccinated elderly mice (100 weeks old) with Ad5.S1 prime−rS1Beta boost had high titers of anti-S1 antibodies after 40 weeks post-boost compared to PBS-immunized mice. We also assessed the effects of a second booster with the recombinant S1 protein of the Omicron variant (BA.1) combined with RS09 peptide (TLR4 agonist) (rS1RS09OM). This booster was effective in stimulating S1-specific immune responses and inducing significantly high cross-ACE2-binding inhibition antibodies against SARS-CoV-2, including Omicron variants, which were correlated with viral neutralizing titers. Furthermore, we compared the immunogenicity of the S1 subunit vaccine in different age groups of mice, 8, 60, and 100 weeks old, corresponding to the ages of administration in this study representing young (<20 years old), middle-aged (42-47 years old), and elderly (>70 years old) humans, respectively (26). The findings in this study provide informative insights into strategies of new emerging SARS-CoV-2 booster vaccination in the elderly population previously vaccinated.

## Results

### Evaluation of Long-Term Immunogenicity and Effect of a Second Booster in Aged Mice

In our previous study, we assessed the efficacy of the rS1Beta protein subunit vaccine as a booster in aged mice that had been previously primed with the adenoviral vaccine, evaluating its effects up to week 80 post-initial vaccination, which is 28 weeks post-1^st^ boost (25). We subsequently extended the study to mice nearing the limit of their lifespan to examine the long-term persistence of immunogenicity. To do so, we initially determined the endpoint titers of antigen-specific IgG antibodies in the sera of vaccinated mice at week 92 post-prime, which corresponded to 40 weeks after the first booster vaccination **(Fig. 1B)**. As shown in Fig. 2A, we observed significantly elevated titers of anti-S1WU IgG antibodies in G4 (Ad5.S1 SC-rS1Beta prime-boost) (*p* < 0.05) even after 40 weeks post-boost, in comparison to the control group treated with PBS group (G1), while comparable titers of anti-S1WU IgG antibodies were present in G5 (Ad5.S1 IN-rS1Beta prime-boost) (*p* = 0.1355).

**Figure 1.**
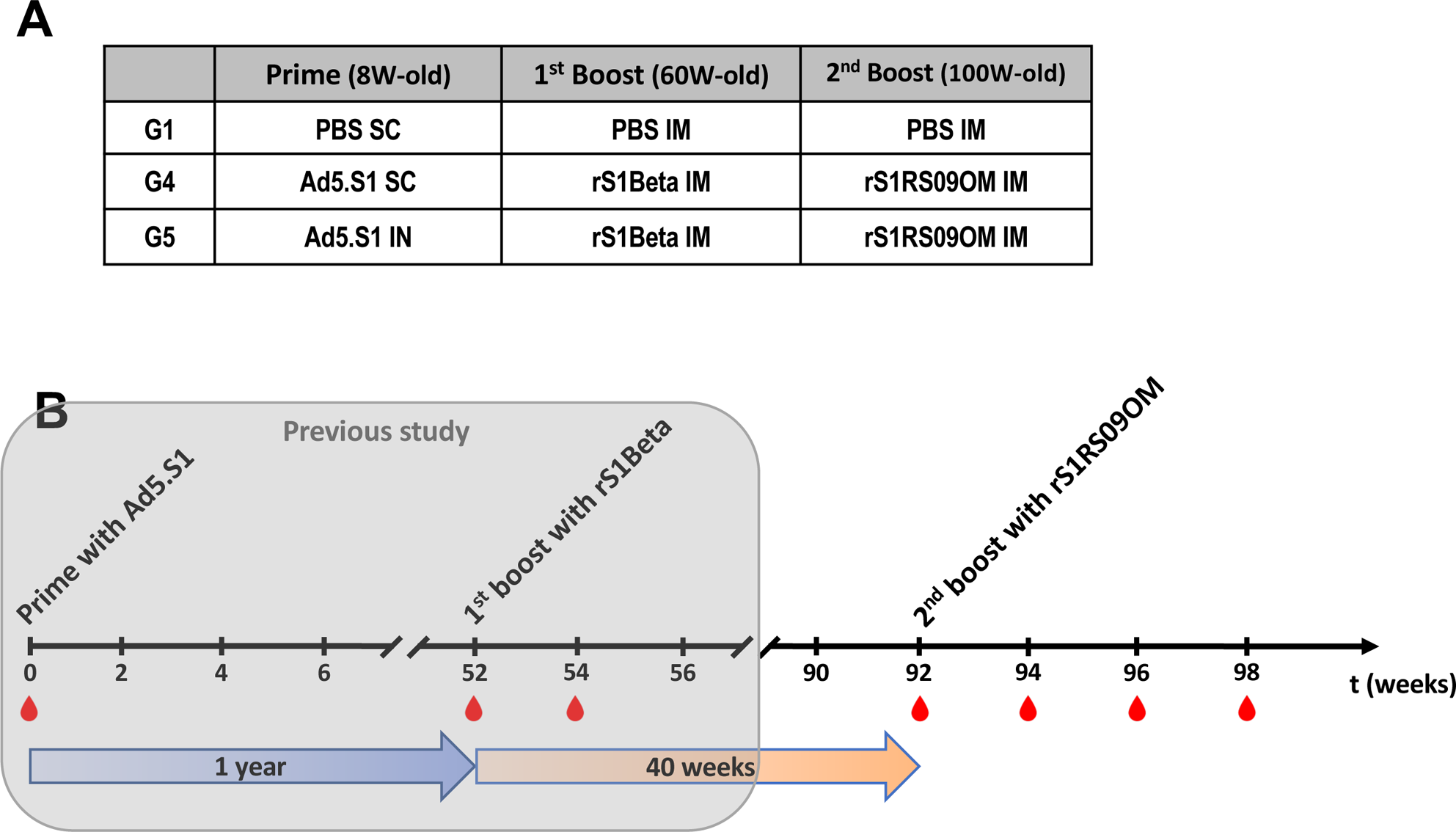
Secondary boost immunization experiments with rS1RS09OM. **(A)** Immunogens and delivery routes of each group **(B)** Schedule of immunization and blood sampling. BALB/c mice were initially primed with 1.5×10^10^ vp of adenoviral vaccine (Ad5.SARS-CoV-2-S1 (Ad5.S1)) SC or IN at 8 weeks old. A negative control group received PBS. One year later, at 60 weeks old, mice were boosted with 15 μg of SARS-CoV-2 rS1Beta recombinant proteins IM. At 40 weeks after the first boost (at week 92 post-prime, 100 weeks old), mice received another boost with 15 μg of rS1RS09OM recombinant proteins IM. The red drops represent the times of blood sampling, and the numbers indicate the weeks after the initial prime immunization.

**Figure 2.**
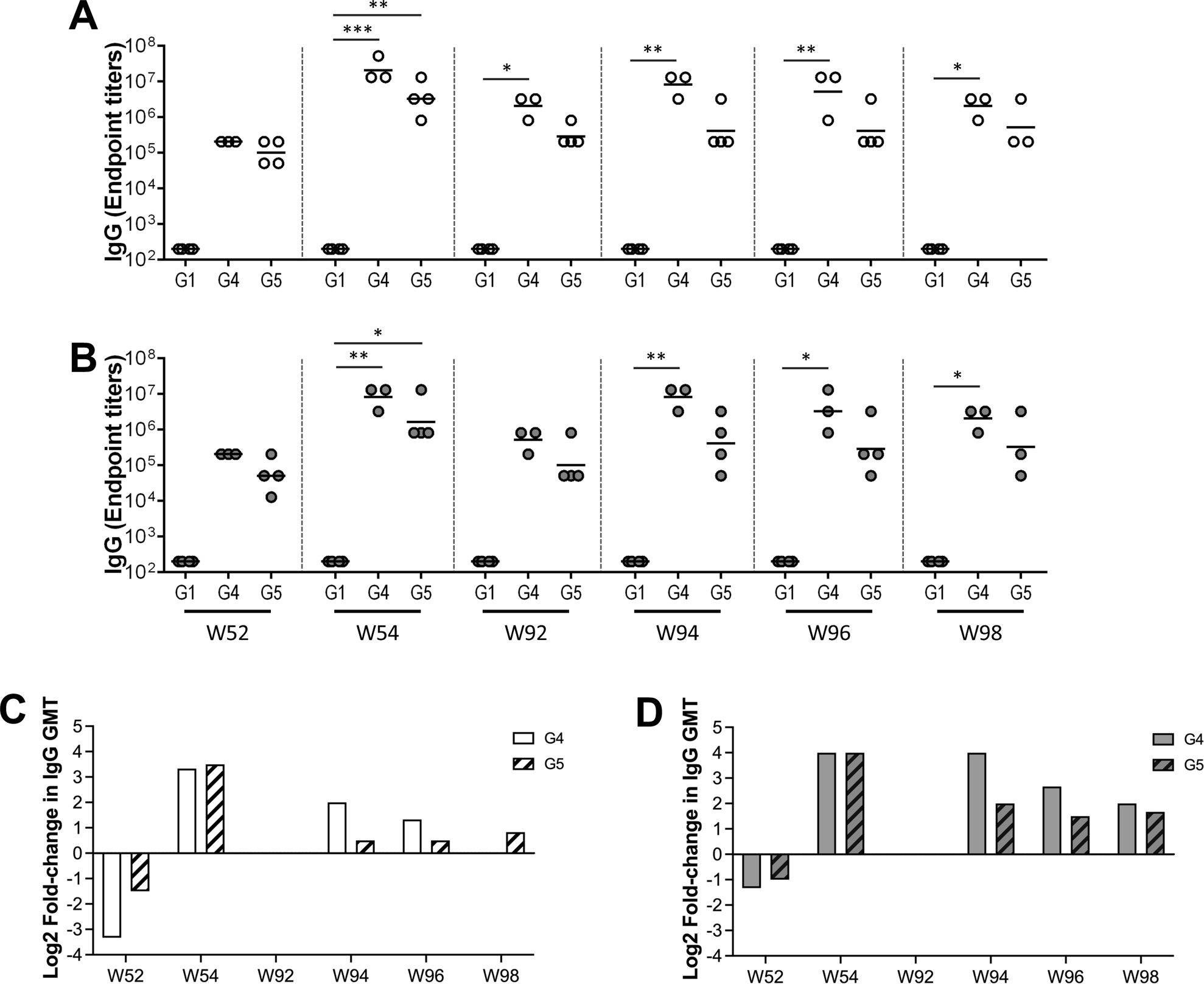
Antigen-specific antibody responses in aged BALB/c mice primed with Ad5.S1 and serially boosted with SARS-CoV-2 rS1Beta and rS1RS09OM protein subunit vaccine. The animal experiment was performed as described in Figure 1. Immune responses were evaluated at weeks 52, 54, 92, 94, 96, and 98 post-prime (N=3 or 4 per group). Reciprocal serum endpoint dilutions of SARS-CoV-2 rS1WU-specific **(A)** and rS1RS09OM-specific **(B)** antibodies were measured using ELISA to determine the IgG endpoint titers. **(C and D)** Fold change of IgG GMT for SARS-CoV-2 rS1WU-specific (white box) and rS1RS09OM-specific (grey box) antibodies in G4 (solid box) and G5 (striped box) after both booster injections relative to those before the second boost (at week 92 post-prime). The horizontal lines represent the geometric mean antibody titers (GMT). Statistical significance was determined using the Kruskal-Wallis test, followed by Dunn’s multiple comparisons (\**p* < 0.05, ∗∗ p < 0.01). The asterisks in Figure 2 indicate statistical differences when compared with G1, the PBS group.

To assess the effect of a second booster with the Omicron recombinant protein S1 subunit vaccine in near lifespan-aged mice, animals were immunized with 15 μg of rS1RS09OM intramuscularly at weeks 92 (100 weeks old) post-prime, and sera were collected at weeks 2, 4, and 6 post-2^nd^ boost (**Fig. 1B**). IgG endpoint titers against both Wuhan S1 (**Fig. 2A)** and Omicron S1 (**Fig. 2B**) were assessed using ELISA. Regarding Wuhan S1, the G4 mouse group exhibited significantly higher geometric mean titers (GMT) at weeks 0, 2, 4, and 6 post-2^nd^ boost when compared to G1. In contrast, the G5 mouse group did not display a significant increase in GMT at any time point. Notably, only one mouse out of the four mice in the G5 group showed an increased GMT after the second boost (**Fig. 2A)**. The fold changes in GMT of the IgG endpoint titers, compared to those at week 92, were 4-, 2.5-, and 1-fold in G4 at week 2, 4, and 6 post-boost, respectively, while it remained relatively stable at 1.4-fold in G5 in all time points (**Fig. 2C**). The fold change in GMT of IgG endpoint titers in G4 and G5 at week 94 was lower than the levels at week 54 (4- and 1.4-fold vs. 10- and 11.3-fold, respectively). Interestingly, the fold change of IgG GMT in G4 and G5 at week 2 after the second boost was lower than that observed after the first boost when compared to levels before each boost (4- and 1.4-fold vs. 101.6- and 32-fold, respectively).

For the Omicron S1, the G4 mouse group had significantly higher GMT at weeks 2, 4, and 6 post-2^nd^ boost compared to the PBS group, while the G5 mouse group had no significant increase in GMT at any time points. However, three out of the four mice in the G5 group showed an increased GMT after the 2^nd^ boost (**Fig. 2B**). The fold change of GMT of the IgG endpoint titer, compared to those at week 92, were 16-, 6.35-, and 4-fold in G4 at weeks 2, 4, and 6 post-2^nd^ boost, respectively, while it was 4-, 2.8-, and 3.1-fold in G5 (**Fig. 2D**). The GMT fold change in IgG endpoint titer for G4 at week 94 was similar with the levels at week 54 (16-fold), while those of G5 at week 94 showed a lower fold change compared those at week 54 (4-fold vs. 16-fold, respectively). Likewise, the fold change of IgG GMT in G4 and G5 at week 2 after the second boost was lower than that observed after the first boost when compared to levels before each boost (16- and 4-fold vs. 40.3- and 32-fold, respectively). In summary, these findings demonstrate that antibody recalls against Wuhan and Omicron S1 were rapid at week 2 following the second booster vaccination with rS1RS09OM subunit vaccine in both G4 and G5 groups. The GMT fold change of IgG endpoint titers of the G4 group waned rapidly, while it remained relatively stable in G5 up to week 6 post-boost. Furthermore, the decline in IgG antibody responses in G4 after a 2^nd^ booster at 100 weeks of age was faster than after the first booster at 60 weeks of age (25). These results suggested that the immune response increases modestly and wanes quickly in elderly mice near the end of their lifespan, even though the recalls are rapid.

### IgG Subtypes Level after a Second Booster

To evaluate the specificity of the IgG antibody response, whether it was skewed towards Th1- or Th2-biased response, we collected serum samples at weeks 0 and 2 after a second booster vaccination (at weeks 92 and 94 post-prime). These samples were serially diluted to determine endpoint titers of rS1WU- and rS1RS09OM-specific IgG1 and IgG2a antibodies for each immunization group, indicating a Th2- or Th1-biased response, respectively (**Fig. 3**). Additionally, to compare IgG1 and IgG2a levels after the first and second boost, serum samples collected at weeks 0 and 2 following the first booster vaccination (at weeks 52 and 54 post-prime) were also analyzed.

**Figure 3.**
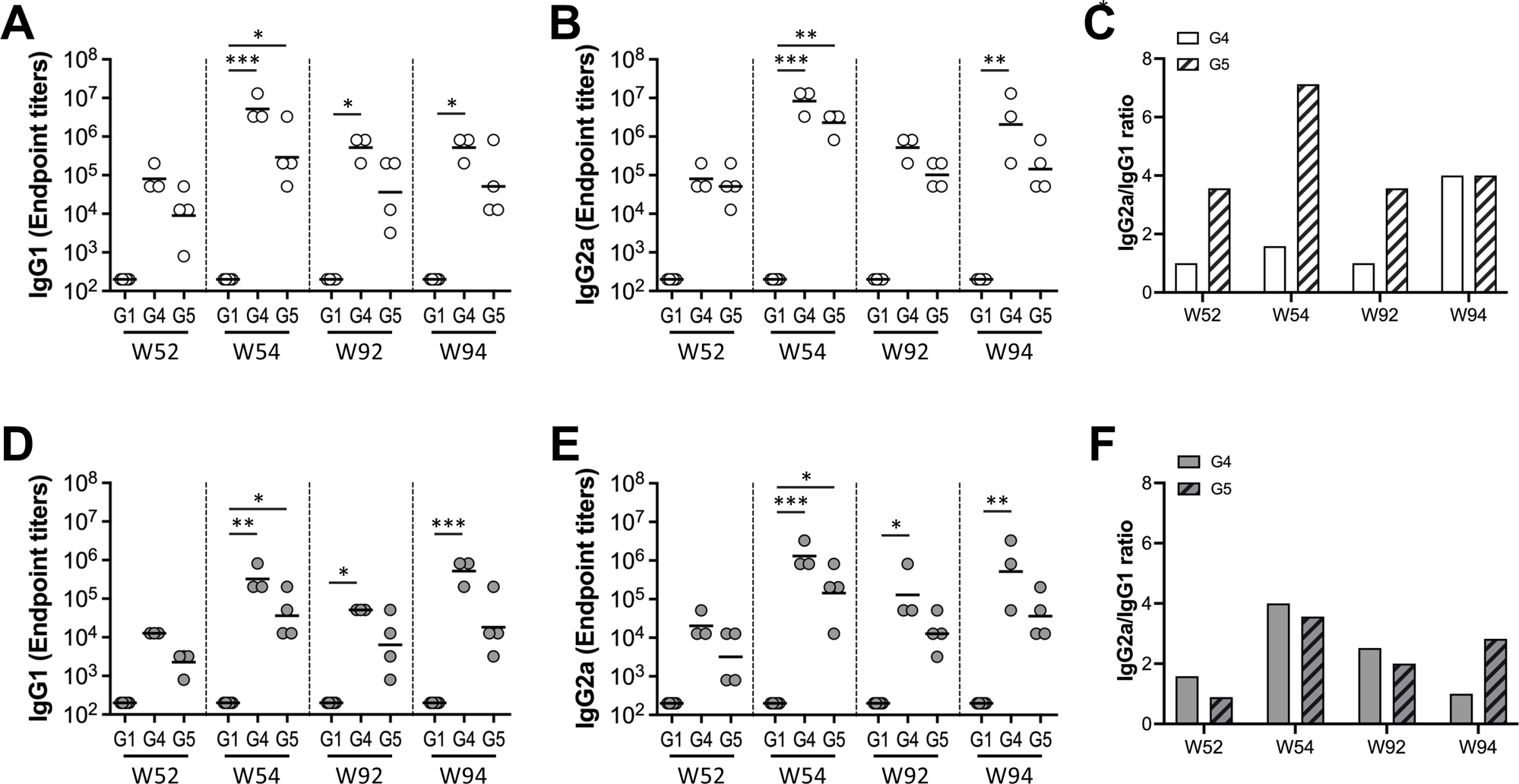
IgG subclasses after both boosters with rS1Beta and rS1RS09OM protein subunit vaccine. Immune responses were assessed at weeks 52, 54, 92, and 94 post-prime, which are before and after each booster (N=3 or 4 per group). Reciprocal serum endpoint dilutions of rS1WU-specific **(A and B)** and rS1RS09OM-specific **(D and E)** antibodies were measured by ELISA to determine the IgG1 **(A and D)** and IgG2a **(B and E)** endpoint titers. The horizontal lines represent the geometric mean antibody titers (GMT). The IgG2a/IgG1 ratio for rS1WU-specific **(C)** and rS1RS09OM-specific **(F)** antibodies was also calculated. Statistical significance was determined using the Kruskal-Wallis test, followed by Dunn’s multiple comparisons (\**p* < 0.05, ∗∗p < 0.01, ∗∗∗p < 0.001). The asterisks in Figure 3 indicate statistical differences when compared with G1, the PBS group.

The induction of rS1WU-specific IgG1 antibodies was significantly higher in G4 at weeks 54, 92, and 94 compared to G1. As shown in Fig. 3A, G4 showed significantly elevated titers of rS1WU-specific IgG1 antibodies after a first boost with rS1Beta at 60 weeks old (*p* = 0.0003 at week 54). However, these levels did not increase following the second boost with rS1RS09OM at 100 weeks old (*p* = 0.0129 at weeks 92 and 94). The rS1WU-specific IgG2a antibodies were significantly induced in G4 at weeks 54 and 94 following both booster vaccinations compared to G1 (*p* = 0.0005 at week 54 and *p* = 0.0074 at week 94). In G5, the induction of rS1WU-specific IgG1 and IgG2a antibodies was only significant at weeks 54 compared to G1 (*p* = 0.0243 and *p* = 0.0015, respectively), while those remained similar following the second boost (**Fig. 3A and 3B**). For the Omicron S1, G4 showed significantly higher rS1RS09OM-specific IgG1 and IgG2a antibodies at weeks 54, 92, and 94 compared to G1. Notably, G4 displayed significantly increased rS1RS09OM-specific IgG1 antibodies following both the first and second boosts (**Fig. 3D and 3E**). In G5, S1RS09OM-specific IgG1 and IgG2a antibodies were significantly increased at week 54 after the first boost at 60 weeks old, with a modest increase following the second boost at 100 weeks old.

Th1-dominant responses were observed in both G4 and G5 at all time points. Notably, the GMT of rS1RS09OM-specific IgG1 at week 94 was higher than that at week 54, while the GMT of rS1WU-specific IgG1 and IgG2a, as well as rS1RS09OM-specific IgG2a, were highest at week 54 after the first boost. Furthermore, the rS1WU-specific IgG2a/IgG1 ratio in G5 was higher than that in G4 at weeks 52, 54, and 92, before the second boost, while rS1RS09OM-specific IgG2a/IgG1 ratio in G5 was higher than that in G4 at weeks 94, after the second boost (**Fig. 3C and 3F**). Together, these findings indicate that a second booster can elicit robust and well-balanced S1-specific antibody responses in mice primed with Ad5.S1 via SC or IN administration at a young age, even in aged mice nearing the end of their lifespan.

### Neutralizing Antibody Levels After a Second Booster

To assess the presence of SARS-CoV-2-specific neutralizing antibodies following a second booster, we conducted a microneutralization assay (VNT_90_). This assay evaluated the ability of sera from immunized mice to neutralize different SARS-CoV-2 variants, including Wuhan, Delta (B.1.617.2), and Omicron (BA.1) variants. After the second booster vaccination, G4 exhibited statistically significant neutralizing activity against all three SARS-CoV-2 variants at week 2 post-2^nd^ boost (at week 94 post-prime) compared to the control group (G1), highlighting the similar neutralizing activity against both Deta and Omicron variants. In contrast, G5 did not show significant differences (*p* = 0.3615 in Wuhan, *p* = 0.0850 in Delta, *p* > 0.9999 in Omicron). Notably, two out of four mice in G5 were shown neutralization activity against Omicron at week 2 following the second boost. The GMT of VNT_90_ in G4 and G5 were as follows: 160.0 and 26.8 against Wuhan, 63.5 and 28.3 against Delta (B.1.617.2), and 57.7 and 8.9 against Omicron (BA.1) at week 2 post-2^nd^ boost, respectively (**Fig. 4**). These findings demonstrate that elderly mice primed with Ad5.S1 subcutaneously (G4) can produce statistically significant neutralizing antibodies against various SARS-CoV-2 variants, including Omicron (BA.1), with similar levels following a second boost with Omicron rS1. In contrast, intranasal priming (G5) fails to induce sufficient neutralizing antibodies against Omicron, although a comparable level of neutralization against Wuahan and Delta variants was shown.

**Figure 4.**
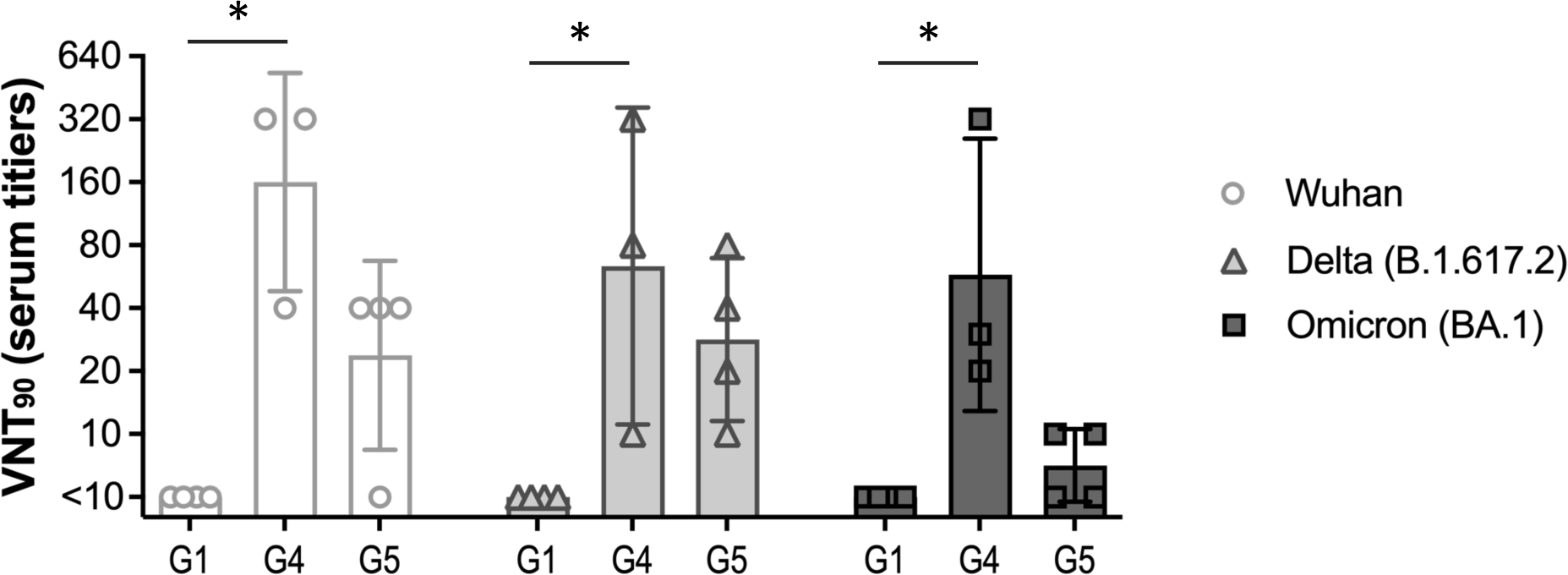
Neutralization of SARS-CoV-2 variants. Sera from mice, after the second boost with rS1RS09OM (at week 94 post-prime), were assessed using a microneutralization assay (NT_90_) to measure neutralization against SARS-CoV-2 variants, including Wuhan (white circle), Delta variant (B.1.617.2, light grey triangle), and Omicron variant (BA.1, dark grey square). The data are presented as geometric means with error bars representing geometric standard deviations (SD). Groups for each variant were compared by the Kruskal-Wallis test, followed by Dunn’s multiple comparisons. Significant differences are indicated by *p < 0.05.

### Boost with rS1RS09OM subunit vaccine induced ACE2 Binding Inhibiting Antibodies against Omicron Variants

We conducted additional tests to assess the ability of serum antibodies to inhibit the interaction between ACE2 and the trimeric spike protein of SARS-CoV-2 variants, representing a sensitive measure of neutralizing activity. We used MSD V-PLEX SARS-CoV-2 (ACE2) Kit Panel 25, which includes Wuhan, Omicron (BA.1), Omicron sub-variants (BA.2, BA.3, BA.1+R346K, BA.1+L452R), Delta lineage (AY.4), Alpha (B.1.1.7), Beta (B.1.351), and France (B.1.640.2) (**Fig. 5**). We examined the sera from all animals in G4 **(Fig. 5A)** and G5 **(Fig. 5B)** at weeks 0, 52, 54, 92, and 94 post-prime at a 1:25 dilution. The ACE2-binding inhibitions of G4 sera at weeks 54 and 94 were significantly increased for all tested SARS-CoV-2 variants, including Omicron and its subvariants when compared to week 0 (**Fig. 5A**). In G5, antibodies showed significant increases only at week 54 compared to week 0 for Wuhan, Delta, Alpha, Beta, France, and Omicron (BA.1+L452R). They demonstrated moderate ACE2-binding inhibition for most Omicron variants (BA.1, BA.2, BA.3, and BA.1+R346K spikes), which were not statistically significant when compared to week 0 (**Fig. 5B**). The increase and decrease in percent inhibition towards the different variants followed the same trend for both groups. Interestingly, the fold change in mean ACE2-binding inhibition at week 94 compared to week 92 was higher for Omicron variants (1.75- to 5.15-fold) than for other parental variants (1.02- to 1.27-fold) in both G4 and G5 (**Fig. 5C**). Specifically, the fold change of ACE2 inhibitory activities for G4 averaged 3.036 ± 0.868 for Omicron variants and 1.092 ± 0.062 for parental variants, respectively. For G5, the averages were 2.912 ± 1.314 and 1.176 ± 0.060, respectively.

**Figure 5.**
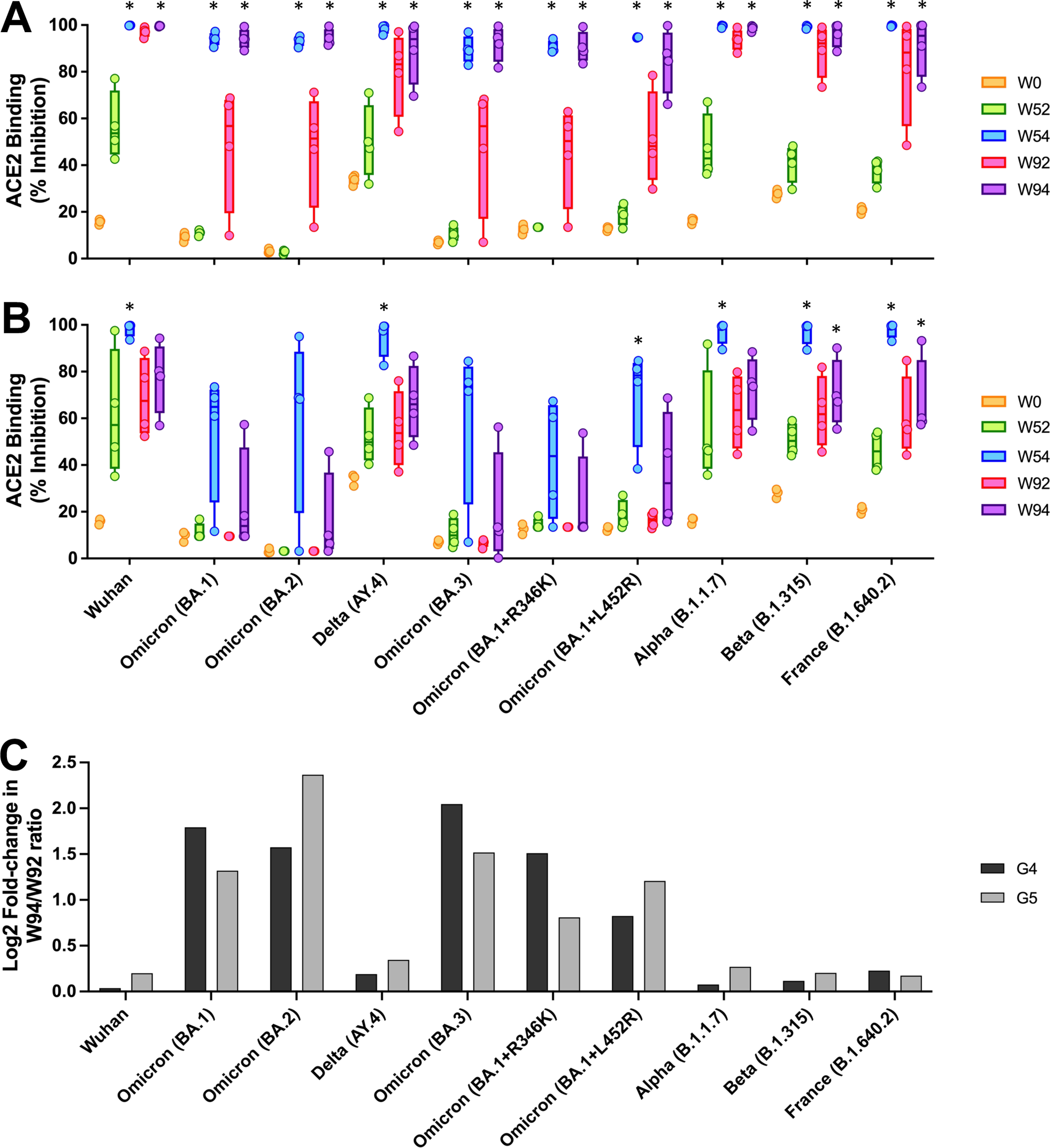
Percent ACE binding inhibition of neutralizing antibodies against SARS-CoV-2 variants. Antibodies in sera capable of neutralizing the interaction between SARS-CoV-2 Wuhan, Omicron (BA.1), Omicron sub-variants (BA.2, BA.3, BA.1+R346K, BA.1+L452R), Delta lineage (AY.4), Alpha (B.1.1.7), Beta (B.1.351), and France (B.1.640.2) variants spike and ACE2 were examined from G4 **(A)** and G5 **(B)** at week 0 (peach), 52 (green), 54 (blue), 92 (pink), and 94 (purple) post-prime. Serum samples were diluted at 1:25 before adding to the V-PLEX plates. Box and whisker plots represent the median and upper and lower quartile (box) with minimum and maximum values (whiskers). Asterisks represent statistical differences compared with pre-immunized sera. **(C)** Fold change of ACE2 binding inhibition (%) in G4 (black box) and G5 (grey box) at week 2 after the second booster injection (at week 94 post-prime) relative to those before the second boost (at week 92 post-prime). Statistical significance was determined using the Kruskal-Wallis test, followed by Dunn’s multiple comparisons (\**p* < 0.05). The asterisks in Figure 5 represent statistical differences when compared with G1, the PBS group.

To further investigate boost-induced neutralizing activities against Omicron variants in G4 and G5 sera, mouse sera were diluted to 1:100 (**Supplementary Fig. 1**). ACE2-binding inhibition significantly increased in G4 sera at weeks 54 and 94 for all SARS-CoV-2 variants, including Omicron sub-lineage variants at a 1:100 dilution. In contrast, G5 sera showed very low ACE2-binding inhibition, with significant increases observed only for the spikes of Wuhan, Delta, Alpha, Beta, and France at week 54 compared to week 0. The fold change in ACE2 inhibitory activities for G4 averaged 2.368 ± 1.086 for Omicron variants and 0.456 ± 0.109 for parental variants. For G5, the averages were 0.636 ± 0.249 and 0.316 ± 0.138, respectively.

### Correlations between the levels of ACE2 inhibition Levels and neutralizing antibodies

Following the administration of a second booster, ACE2 binding inhibition and VNT_90_ titers increased significantly against Wuhan, Delta (B.1.617.2), and Omicron (BA.1) compared to pre-vaccinated sera, with no significant differences found among the variants. To determine the correlations between ACE2 inhibition levels and neutralizing antibodies, we conducted correlation analyses on ACE2 inhibition using 1:25 diluted mouse sera and log-transformed VNT_90_ data of Wuhan, Delta (B.1.617.2), and Omicron (BA.1) at week 2 post-second booster. A positive correlation between V-PLEX ACE2 inhibition and VNT_90_ was observed in all animals from G1, G4 and G5 at week 2 post-second boost (Spearman’s correlation coefficients, *r* = 0.9163 (95% CI: 0.8264-0.9606, *p* <0.0001) (**Fig. 6A**). The Spearman’s correlation coefficient was slightly lower when the analysis was performed with 1:100 diluted mouse sera (*r* = 0.8827 (95% CI: 0.7615-0.9163, *p* <0.0001) (**Supplementary Fig. 2A**).

**Figure 6.**
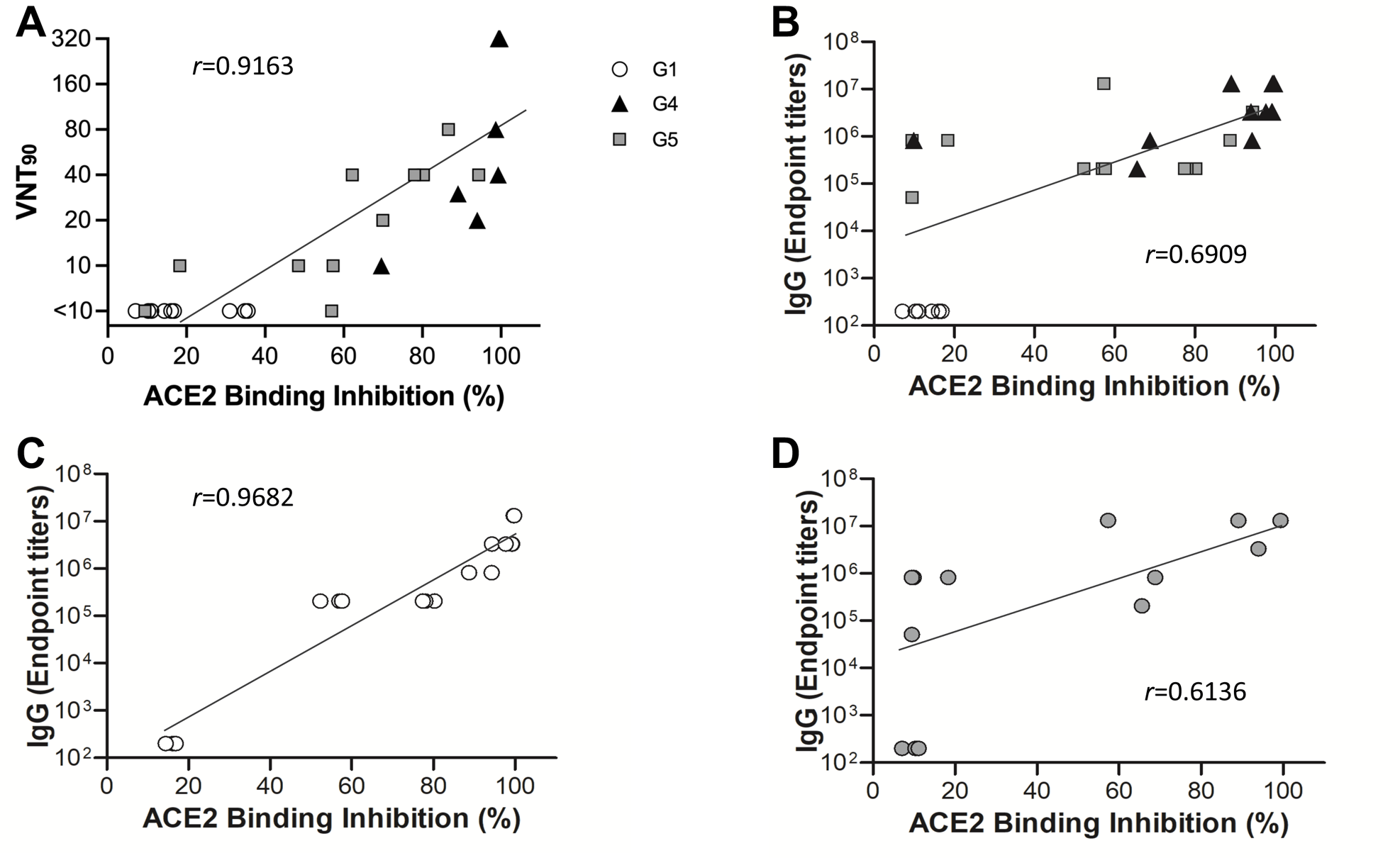
Correlation betseen the VNT_90_ or SARS-CoV-2-S1-specific IgG titers and ACE2 binding inhibitions. **(A)** Correlation between VNT_90_ (Log_2_) against Wuhan, Delta (B.1.617.2), and Omicron (BA.1), and ACE2 binding inhibition (%) of 1:25 diluted sera from all animals of G1 (white circle), G4 (black triangle), and G5 (Grey square) at week 94 (**B)** Correlation between SARS-CoV-2-S1-specific IgG titers against Wuhan and Omicron (B.1.617.2), and ACE2 binding inhibition (%) of 1:25 diluted sera from all animals of G1 (white circle), G4 (black triangle), and G5 (grey square) at weeks 92 and 94. Correlation between SARS-CoV-2 S1WU-specific (white circle) (**C**) or S1OM-specific (grey circle) (**D**) IgG titer, and ACE2 binding inhibition (%) of 1:25 diluted sera among sera from all animals of B. The lines represent the regression line of all samples and each symbol represents an individual mouse. Correlation analysis and calculation of Spearman’s correlation coefficients was performed using GraphPad Prism v9.

Furthermore, a positive correlation between V-PLEX ACE2 inhibition and IgG EPT was observed in all animals from G1, G4, and G5 at weeks 0 and 2 post-second boost (Spearman’s correlation coefficients, *r* = 0.6909 (95% CI: 0.4519-0.8373, *p* <0.0001) (**Fig. 6B**). There was a higher correlation between ACE2 binding inhibition against Wuhan spike and rS1WU-specific IgG EPT (Spearman’s correlation coefficients, *r* = 0.9682 (95% CI: 0.9094-0.9891, *p* <0.0001) when compared to the counterparts against Omicron spike and rS1OM-specific IgG EPT (Spearman’s correlation coefficients, *r* = 0.6136 (95% CI: 0.1734-0.8494, *p* = 0.0088) (**Fig. 6C and D**). The Spearman’s correlation coefficient was slightly higher when the analysis was performed with 1:100 diluted mouse sera (*r* = 0.7194 (95% CI: 0.4959-0.8538, *p* <0.0001) (**Supplementary Fig. 2B**). Likewise, there was a stronger correlation between ACE2 binding inhibition and rS1WU-specific IgG EPT (Spearman’s correlation coefficients, *r* = 0.9682 (95% CI: 0.9094-0.9891, *p* <0.0001) compared to rS1OM-specific IgG EPT (Spearman’s correlation coefficients, *r* = 0.7065 (95% CI: 0.3281-0.8895, *p* = 0.0015) (**Supplementary Fig. 2C and 2D**). Taken together, a single dose of non-adjuvanted recombinant Omicron S1 protein subunit vaccine, when used as a second booster in elderly mice previously prime-boosted with Ad5.S1-rS1Beta, induced broadly cross-reactive neutralizing antibodies against a wide range of SARS-CoV-2 variants, including Omicron and its subvariants, which was correlated with the inhibition of spike-ACE2 binding and IgG levels.

### Systemic Antibody Responses Decrease with Age

The study found that systemic antibody responses decline with age. This decline was particularly evident when evaluating the effect of a boost with rS1RS09OM in the elderly mice (100 weeks old) compared to middle-aged mice (60 weeks old). Therefore, we assessed the humoral immune response in mice of 8, 60, and 100 weeks old to determine the impact of aging on immunogenicity. Young adult BALB/c mice (8 weeks old, n =5), middle-aged mice (60 weeks old, n=5) and elderly mice (100 weeks old, n=4) were intramuscularly primed with 15 μg of SARS-CoV-2 rS1WU or rS1RS09OM, respectively, and received a homologous booster at week 3 post-priming (**Fig. 7A**). Serum samples were collected at weeks, 0, 3, 5, and 7 post-priming and were serially diluted to determine antigen-specific IgG titers. Young mice were evaluated for IgG titer against Wuhan S1 (S1WU), while middle-aged and elderly mice were assessed for Omicron S1 (rS1RS09OM) for, respectively. At 3 weeks after priming, no or very low levels of S1-specific IgG antibodies were observed in all groups of mice. However, at weeks 5 and 7, a significantly high level of S1-specific IgG antibodies was induced in the young mice group (*p* = 0.0070), but not in the middle-aged and elderly mice groups compared to their levels at week 0 (**Fig. 7B**). The mean IgG endpoint titer changes were decreased by 3.2-fold at week 5 and 6.4-fold at week 7 in 60-week-old group compared to the young mice group, while the reductions were even more substantial in the 100-week-old group with 6.1-fold and 23.4-fold, respectively. (**Fig. 7C**). The serum IgG1 responses were statistically significant in the young mice group post-boost, while no significant differences were observed in the middle-aged and elderly mice groups as they aged. On the other hand, the serum IgG2a responses were weak even in the young mice group, and they were negligible in the middle-aged and elderly mice groups (**Fig. 7D and E**). These results suggest that the immune response diminishes with age and shows more pronounced defects in inducing IgG2a (Th1) compared to IgG1 (Th2) isotype antibodies, especially when compared to the responses observed in the young mice group.

**Figure 7.**
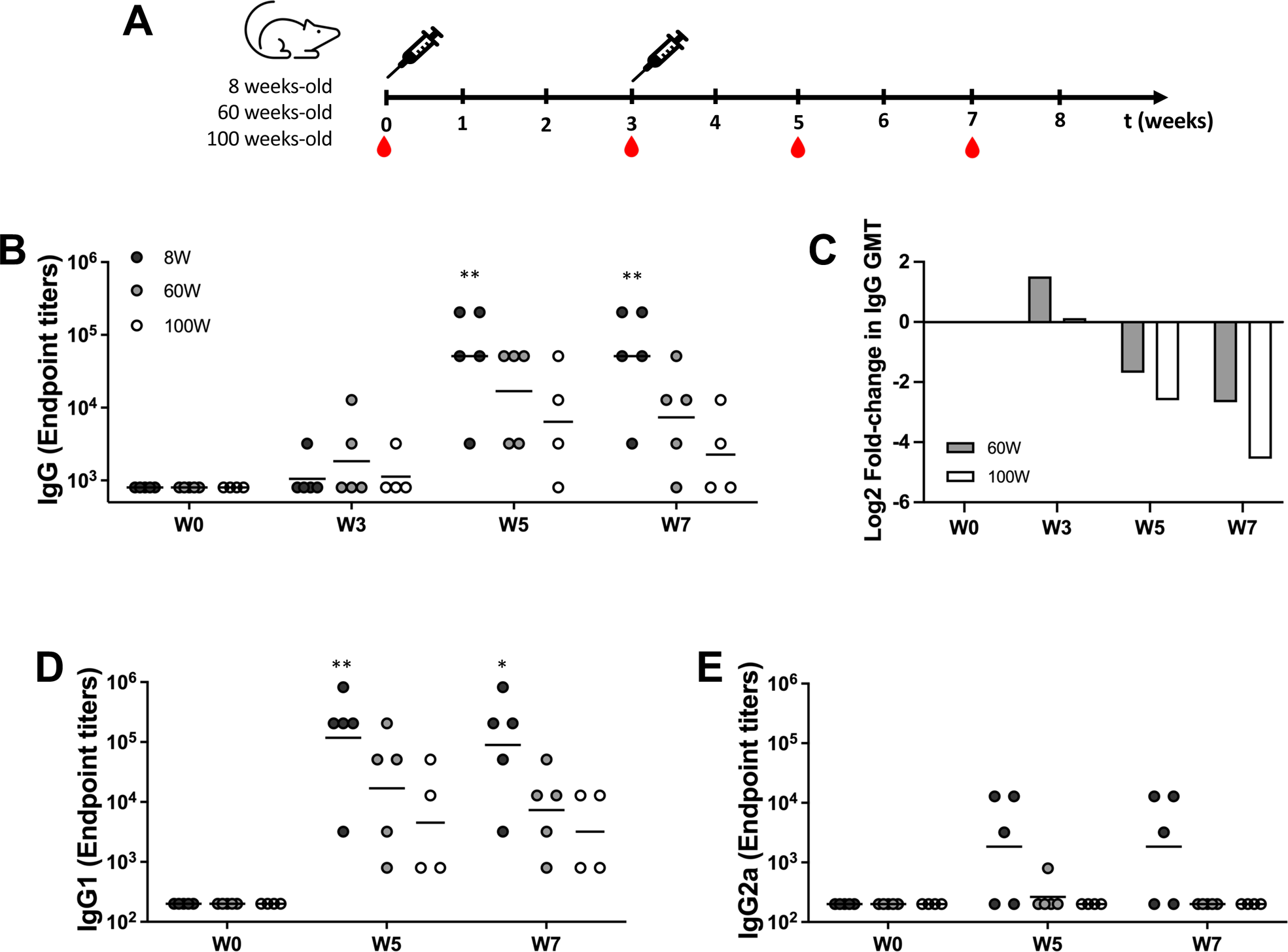
Caparison of immunogenicity of rS1 subunit vaccine in young, middle-aged, and elderly mice. **(A)** Schedule of immunization and blood sampling. 8-, 60-, and 100-week-old BALB/c mice were primed IM with 15 μg of SARS-CoV-2 rS1WU or rS1RS09OM, respectively, and received a homologous booster at week 3 post-prime. The red drops indicate the times of blood sampling. **(B)** The immune responses were assessed at weeks 0, 3, 5, and 7 post-prime (N=4 or 5 per group). Reciprocal serum endpoint dilutions from mice of 8 weeks old (black circle), 60 weeks old (grey circle), and 100 weeks old (white circle) of SARS-CoV-2 rS1WU-specific or rS1RS09OM-specific antibodies were measured by ELISA to determine the IgG endpoint titers. **(C)** Fold change of IgG GMT of middle-aged (grey) and elderly (white) mice relative to those of young mice at each time points. IgG1 **(D)** and IgG2a **(E)** endpoint titers at weeks 0, 5, and 7 post-boost for each age group. Statistical significance was determined by the Kruskal-Wallis test, followed by Dunn’s multiple comparisons (\**p* < 0.05). Asterisks represented statistical differences when compared with pre-immunized sera.

## Discussion

We previously reported that a booster of non-adjuvanted recombinant S1 protein of Beta variant to the aged BALB/c mice immunized with our adenovirus-based COVID-19 vaccine (Ad5.S1) via either I.N. or S.C. induced robust balanced long-lasting IgG antibodies and neutralizing antibodies, which were broadly cross-reacting with SARS-CoV-2 variants (25). In this follow-up study, we demonstrated that a second booster of non-adjuvanted recombinant Omicron S1 protein with TLR4 agonist to the elderly BALB/c from the previous experiment induced a robust balanced humoral immune response, which were broadly cross-reacting with SARS-CoV-2 variants including Omicron and its subvariants and correlated with ACE2-spike interaction inhibition.

The second boost of the Omicron rS1 protein subunit vaccine did not increase as significantly as after the first boost of rS1Beta in mice primed with Ad5.S1 (**Fig.2**). It can be explained by three aspects. One is the age of the mice at the time of boost immunization. Indeed, 21-month-old mice boosted with mRNA elicited lower levels of neutralizing antibodies against Omicron compared to 13-month-old mice (27). These results were parallel to previous findings that individuals 70 years or older (median age 73, range 70-75) who received a primary two-dose schedule with AZD1222 and booster third dose with mRNA vaccine achieved significantly lower neutralizing antibody responses against SARS-CoV-2 spike pseudotyped virus compared with those younger than 70 (median age 66, range 54-69) at 1 month post booster (28). The second reason may be that the vaccine had already reached a high level, and further increases were limited. This phenomenon might be attributed to the loss of short-lived plasma cells in the early stages, causing the initial rapid decline in S1-specific antibodies during the prime-boost. Conversely, the development of long-lived memory plasma cells with additional boosts could explain the plateau in the antibody response (29). The relatively modest increase in IgG levels between the second and third doses could be attributed to the short interval, which might not provide sufficient time for the IgG titer to decline. The last reason may be the antigen from Omicron variants, which is drifted antigenically far from the ancestral Wuhan strain, resulting in a distant location on the phylogenetic tree (30). Likewise, the early nonhuman primate studies suggest that Omicron-specific boosters offer no advantage over a third dose of current vaccines (31). Moreover, preliminary data from small preclinical studies suggested that Omicron-specific vaccines did not perform better than the original vaccines (32). As a consequence, the subunit boost vaccine of ancestral spike provides greater protection in the airways, with superior and better viral control in the upper respiratory airways compared to controls and BA.1 spike subunit vaccine in nonhuman primates (24). Although the BA.1 spike subunit vaccine following the primary series of mRNA-1273 recalled a greater proportion of BA.1 to WA-1-specific memory B cells compared to the ancestral spike-specific vaccine, a large portion of BA.1 response was targeted outside of RBD (24). Hence, a second bivalent boost shot will have the potential benefit to elderly people to induce a strong immune response to protect from reinfection of new emerging SARS-CoV-2 variants. That’s why the U.S. Food and Drug Administration (FDA) and Centers for Disease Control and Prevention (CDC) were offering a second bivalent booster shot to 65 and older people and those with weak immune systems (33).

Although there were no significant differences between mice groups primed with Ad5.S1 SC or IN, animal group primed SC showed a higher GMT of IgG, IgG1, IgG2a, and neutralization antibodies after the second booster than animal group primed IN **(Fig. 2, 3, and 4)**. Other studies showed that heterologous vaccines elicit strong and long-lasting immune responses in the bloodstream as well as promoting elevated IgA levels in the mucosal regions when administered via mucosal routes (34–37). Thus, the mucosal immunity triggered by IN vaccination might be beneficial in the clearance of invading virus, leading to a promising preventive strategy for SARS-CoV-2 infection. The second boost vaccination with subunit vaccine elicited both high S1-specific IgG1 and IgG2a subclass antibodies in elderly mice prime-boosted with Ad5.S1-rS1Beta, indicating a balanced Th1/Th2 response (**Fig. 3**), whereas subunit vaccine alone induced high IgG1 with lower IgG2a leading a possibility of vaccine-associated enhanced respiratory disease (VARED) (38) (**Fig. 7**). Especially, prime-boost vaccination with the protein subunit vaccine in 60- and 100-weeks old mice induced under the limit of detection of IgG2a antibodies (**Fig. 7E**), which play an important role for antiviral immune response though antibody-dependent cellular phagocytosis (ADCP) and antibody-dependent complement deposition (ADCD) (15). Moreover, the homologous Adenovirus-based boost was not effective due to pre-existing neutralizing antibodies against adenovirus. Indeed, Adenoviral-vectored vaccines were less effective than mRNA-based or subunit vaccines as a boost following CoronaVac/AZD1222 prime (19, 39). Furthermore, many recent studies showed spike subunit booster vaccines (CoV2 preS dTM-AS03, NVX-COV2373/NVX-COV2515, or MVC-COV1901) elicited robust neutralizing antibodies against multiple variants in populations who were previously primed with adenovirus-vectored vaccines such as Ad26.CoV2.S or ChAdOx1nCoV-19, although mRNA booster vaccine enhanced neutralization capacity against Omicron (7, 18, 19, 22, 23). Thus, our future work will include more investigation in neutralization activity against spikes of currently circulating variants of interest (VOIs) such as, XBB.1.5., XBB.1.16., and EG.5.

Antigen-specific systemic antibody responses were decrease with age related to many infectious diseases such as respiratory syncytial virus infection (RSV) and influenza (40, 41). In this study, we evaluated the immune response at the age of both booster injections of at 60 and 100 weeks old compared to those at 6 weeks old. We injected the mouse with the same amount of unadjuvanted subunit vaccine intramuscularly and assessed S1-specific immune response by ELISA. As shown in **Fig.7**, the IgG levels decreased with age, especially IgG2a representing Th1 biased immune response was under a detectable level at the age of both booster vaccination (**Fig. 7E**). This study highlights the potential benefit of the booster with protein subunit vaccine in elderly mice who had been exposed to antigen early in life, resulting in enhanced cross-protective vaccine efficiency compared to those who did not expose during their youth. Indeed, mice primed with Influenza A nucleoprotein or matrix 2 and boosted with recombinant adenovirus expressing the same antigens (A/NP-rAd or A/M2-rAd) during their youth were partially protected against challenge 16 months later when they were elderly, whereas not in elderly mice vaccinated with same regimens (40). Therefore, vaccination in childhood demonstrates a potential benefit to protect vulnerable older populations, because the robust and long-lasting immune responses can be elicited from memory B cells by a boost at an elderly age. Of note, while increasing age was associated with decreased humoral immune responses in the present study, 60-week-old mice induced differently with or without vaccination of Ad5.S1 during their youth, resulting in 1024- and 194-fold increased IgG GMT in the mice primed with Ad5.S1 SC or IN, respectively, compared to that of in mice primed-boosted with subunit vaccine when they were 60 weeks-old (**Fig.2A and 7B**).

Here we showed the immunogenicity of the non-adjuvated protein subunit vaccine as a prime and booster effect. However, there will be a beneficial effect of an adjuvanted subunit booster strategy for protection, especially for the aged population. AS01-like adjuvanted SARS-CoV-2 spike subunit vaccine enhanced Th1 type-IgG2a isotype, neutralizing antibodies, and IFN-γ secreting T cell immune responses in both young adult and old-aged mice (42). The AH: CpG-adjuvanted RBD subunit vaccine elicited a robust anti-RBD immune response in aged mice, with an additional boost dose generating an anti-RBD antibody response comparable to young adult mice and providing complete protection from live SARS-CoV-2 challenge (43).

While our results show promising immune responses to the second boost with subunit vaccine of the new emerging variant in elderly mice, there are several limitations, which can be addressed in future research. These include T-cell-specific cellular immunity, the mucosal immune response, the SARS-CoV-2 challenge, and the durability of the immune response, to provide further insights into the effectiveness of the vaccine in preventing infection and disease by a second booster vaccination. However, various studies have reported previously that T-cell immunity was activated after a booster (31, 34, 44, 45). Boosting with NVX-CoV2373 (nanoparticle of spike protein with Matrix-M^TM^ adjuvant) to individuals primed with two doses of ChAdOx1nCov-19 or MVC-COV1901 (protein subunit vaccine of stable prefusion spike protein) of individuals primed with two doses of AZD1222 induced T cell-mediated immune response (19, 21). Furthermore, S-specific T-cell responses were positively correlated with the presence of S-specific binding antibodies (44), implying the induction of robust T-cell immune response after rS1RS09OM booster in this study.

Additionally, our study could not evaluate the durability of the antibody response generated by the 2^nd^ boost vaccination over a longer period, because the mice were already near lifespan. Of note, a considerable number of aged mice developed conditions such as frailty and gross tumors over the 23-month study period, and only *N* = 10 out of the initial *N* = 15 could be assessed for immunogenicity of a second boost (*N* = 3 in group 1 and 4 and *N* = 4 in group 5). A longer time course analysis is needed to assess the durability of the enhanced immune responses induced by a booster dose in the future. However, it might persist for a longer period based on our previous study showing that antibody responses after a boost sustained longer time than after a prime (25). Notably, we have previously published a report describing the robust and long-lasting immunogenicity of a boost with S1 protein subunit of MAP platform than a conventional intramuscular injection to mice, suggesting a boost with MAP of S1 subunit vaccine may also be beneficial in the context of S1 based SARS-CoV-2 vaccines efficiency and equality worldwide (46, 47).

Overall, our study evaluated the effect of the 2nd booster with heterologous S1 protein subunit vaccine in elderly mice after priming of adenoviral vaccines as a pre-clinical model of elderly people immunized with the current approved COVID-19 vaccines. Our findings may have implications for further study of using the MAP platform of recombinant protein S1 subunit vaccine as a booster to enhance cross-neutralizing antibodies against new emerging variants of concern and the potential utility of the booster vaccine in an infrastructure-limited environment.

## Materials and methods

### Antigens

The recombinant protein Omicron S1 with TLR4 agonist (RS09) was produced by transient expression in Expi293 cells, as previously reported (25, 48, 49). In brief, rS1RS09OM was produced by transient expression in Expi293 cells with pAd/S1RS09OM, using ExpiFectamie^TM^ 293 Transfection Kit (ThermoFisher) as reported previously (25, 48, 49).

The recombinant proteins were purified using a CaptureSelect^TM^ C-tagXL Affinity Matrix prepacked column (ThermoFisher), followed the manufacturer’s guidelines as reported previously (25, 48, 49). Briefly, the C-tagXL column was conditioned with 10 column volumes (CV) of equilibrate/wash buffer (20 mM Tris, pH 7.4) before sample application. The supernatant was adjusted to 20 mM Tris with 200 mM Tris (pH 7.4) before being loaded onto a 5-mL prepacked column with a 5 ml/min rate. The column was then washed by alternating with 10 CV of equilibrate/wash buffer, 10 CV of strong wash buffer (20 mM Tris, 1 M NaCl, 0.05% Tween-20, pH 7.4), and 5 CV of equilibrate/wash buffer. The recombinant proteins were eluted from the column using elution buffer (20 mM Tris, 2 M MgCl_2_, pH 7.4). The eluted solution was then desalted and concentrated with preservative buffer (PBS) in an Amicon Ultra centrifugal filter device with a 50,000 molecular weight cutoff (Millipore). The concentration of the purified recombinant proteins was determined using the BCA protein assay kit (Thermo Scientific), with bovine serum albumin (BSA) as the protein standard. The proteins were separated by reducing SDS-PAGE and visualized by silver staining.

### Animals and immunization

Female BALB/c mice (n = 5 animals per group) primed with adenovirus-based COVID-19 vaccine (Ad5.S1) SC or IN at 8 weeks old (1) and boosted with 15 μg of rS1Beta (25) IM at week 52 post-priming (60 weeks old). Subsequently, these mice (n = 3 or 4 animals per group) received a secondary boost by IM with 15 μg of rS1RS09OM (49) at week 92 post-priming (100 weeks old) in the thigh or PBS was administered as a negative control. Blood samples were collected via the retro-orbital vein at weeks 0, 2, 4, and 6 after the secondary boost immunization. The obtained serum samples were diluted and used to evaluate both rS1WU- and rS1RS09OM-specific antibodies by enzyme-linked immunosorbent assay (ELISA). Since aged mice develop spontaneous leukemias and other tumors, the dedicated veterinarians oversee the animals’ physical and psychological health and ruled out mouse having disease that may influence immune responses. Indeed, at the age of 100 weeks, only 3 or 4 mice per group survived and others were euthanized due to natural tumors. Data at weeks 0, 52, and 54 are presented for those that were alive at week 92 post-priming.

At week 0, young, middle-aged, or elderly female BALB/c mice [n = 5 (8 weeks old), 5 (60 weeks old), and 4 (100 weeks old) animals per group] were bled from the retro-orbital vein and primed with 15 μg of either rS1WU (8W) or rS1RS09OM (60W and 100W). Mice were bled at week 3 and received a 15 μg homologous booster. Mice were bled at weeks 5 and 7 post-priming. All mice were maintained under specific pathogen-free conditions at the University of Pittsburgh, and all experiments were conducted in accordance with animal use guidelines and protocols approved by the University of Pittsburgh’s Institutional Animal Care and Use Committee (IACUC).

### ELISA

To investigate the immunogenicity of SARS-CoV-2 S1 recombinant proteins, we conducted an enzyme-linked immunosorbent assay (ELISA) following previously established protocols (25, 46). Sera from all mice were collected prior to the second boost (week 92) and every two weeks thereafter (week 94, 96, 98) after immunization. These sera were tested for SARS-CoV-2 S1WU- or S1RS09OM-specific IgG antibodies using conventional ELISA. Additionally, sera collected at week 52, 54, 92, and 94 before and after the first and second boost vaccination were tested for SARS-CoV-2-S1-specific IgG1 and IgG2a antibodies using ELISA. Briefly, ELISA plates were coated with 200 ng of recombinant SARS-CoV-2-S1WU protein (Sino Biological) or SARS-CoV-2-S1RS09OM protein per well and incubated overnight at 4°C in carbonate coating buffer (pH 9.5) and then blocked with PBS containing 0.05% Tween 20 (PBS-T) and 2% bovine serum albumin (BSA) for one hour. Mouse sera were serially diluted in PBS-T with 1% BSA and incubated overnight. After washing the plates, anti-mouse IgG-horseradish peroxidase (HRP) (1:10000, Jackson Immunoresearch) was added to each well and incubated for one hour. For IgG1 and IgG2a, biotin-conjugated IgG1 and IgG2a (1:1000, eBioscience), followed by HRP-conjugated avidin (Av-HRP) (1:50000, Vector Laboratories) were added to each well and incubated for 1 hour. The plates were washed three times and developed with 3,3’5,5’-tetramethylbenzidine, and the reaction was stopped. Absorbance at 450 nm was determined using an ELISA reader (Molecular Devices SPECTRAmax).

### SARS-CoV-2 microneutralization assay

The neutralizing antibody titers against SARS-CoV-2 were determined using the following protocol (50, 51). Initially, 50 µl of sample from each mouse, starting at a 1:10 dilution and progressing in twofold dilution, were added to two wells of a flat-bottom tissue culture microtiter plate (COSTAR). These samples were then mixed with an equal volume of 100 TCID_50_ of a SARS-CoV-2 strain, the Wuhan, Delta (B.1.617.2), or Omicron (BA.1) strain, which had been isolated from symptomatic patients and previously titrated. After a 1-hour incubation at 33°C in a 5% CO_2_ environment, 3 x 10^4^ VERO E6 cells were added to each well. Following 72 hours of incubation at 33°C with 5% CO_2_, the wells were stained with Gram’s crystal violet solution along with 5% formaldehyde (40% m/v, Carlo ErbaSpA, Arese, Italy) for 30 min. After washing, the wells were assessed to evaluate the degree of cytopathic effect (CPE) compared to the virus control. The neutralizing titer was determined as the highest dilution that resulted in a 90% reduction in CPE. A positive titer was considered equal to or greater than 1:10. The GMT of VNT_90_ endpoint titer was calculated with 5 as a negative shown <1:10. Sera from mice before vaccine administration were always included in VNT assay as a negative control.

### ACE2 blocking assay

Antibodies that block the binding of SARS-CoV-2 spike proteins, including Wuhan, Omicron (BA.1), Omicron sub-variants (BA.2, BA.3, BA.1+R346K, BA.1+L452R), Delta lineage (AY.4), Alpha (B.1.1.7), Beta (B.1.351), and France (B.1.640.2), to ACE2 receptors were detected using the V-PLEX SARS-CoV-2 Panel 25 (ACE2) Kit from Meso Scale Discovery (MSD), following the manufacturer’s instructions. The assay plate was blocked for 30 min and then washed. Serum samples were diluted (1:25 or 1:100), and 25 μl of each diluted sample was added to individual wells. The plate was incubated at room temperature for 60 min with shaking at 700 rpm. Following this, SULFO-TAG conjugated ACE2 was added, and incubation continued with shaking for 60 min. The plate was washed, and 150 μl of MSD GOLD Read Buffer B was added to each well. Subsequently, the plate was read using the QuickPlex SQ 120 Imager, and Electrochemiluminescent values (ECL) were generated for each sample. Values lower than those observed in pre-immunized sera were adjusted using the values at week 0. Results were calculated as a percentage inhibition in comparison to the negative control for the ACE2 inhibition assay, and this percentage of inhibition is calculated as follows: % inhibition = 100 × (1 − (sample signal/negative control signal)).

### Statistical analysis

Statistical analyses were performed using GraphPad Prism v9 (San Diego, CA). Antibody endpoint titers and neutralization data were analyzed using the Kruskal-Wallis test, followed by Dunn’s multiple comparisons. Intergroup statistical comparison was performed with a Mann-Whitney U test. Data were presented as either means ± standard errors of the mean (SEM) or the geometric mean ± geometric errors. Significant differences were denoted by asterisks (∗ p < 0.05, ∗∗p < 0.01, ∗∗∗p < 0.001). Correlations between the V-Plex ACE2-blocking and VNT_90_ or IgG endpoint titers were determined using correlation analysis and calculations of Spearman coefficients and 95% confidence intervals (CIs).

## ACKNOWLEDGMENTS

This work was supported by National Institutes of Health grants (UM1-AI106701, R01DK119936-S1, and U01-CA233085) and UPMC Enterprises IPA 25565. The funders had no role in study design, data collection and analysis, decision to publish, or preparation of the manuscript.

We declare that we have competing interests in relation to the research presented in the manuscript. A.G. and E.K. are cofounders of Gaphas Pharmaceutical Inc., a private startup company that may potentially benefit from the findings of this research. A.G., E.K., and M.S.K. have equity in Gaphas Pharmaceutical Inc. However, we have taken measures to ensure that the research is conducted objectively and that the data and conclusions presented in the manuscript are not influenced by our competing interests. The study was designed, conducted, and analyzed independently of the company. We also declare that Gaphas Pharmaceutical Inc. did not provide the financial or material support for this research.

---

**Supplementary Figure 1. Percent ACE binding inhibition of neutralizing antibodies against SARS-CoV-2 variants.** Antibodies in sera capable of neutralizing the interaction between SARS-CoV-2 Wuhan, Omicron (BA.1), Omicron sub-variants (BA.2, BA.3, BA.1+R346K, BA.1+L452R), Delta lineage (AY.4), Alpha (B.1.1.7), Beta (B.1.351), and France (B.1.640.2) variants spike and ACE2 were examined from G4 **(A)** and G5 **(B)** at week 0 (peach), 52 (green), 54 (blue), 92 (pink), and 94 (purple) post-prime. Serum samples were diluted in 1:100 before adding the V-PLEX plates. Box and whisker plots represent the median and upper and lower quartile (box) with min and max (whiskers). Asterisks represent statistical differences compared with pre-immunized sera. **(C)** Fold change of ACE2 binding inhibition (%) in G4 (black box) and G5 (grey box) at week 2 after a 2nd booster injection (at week 94 post-prime) relative to those of pre-2^nd^ boost (at week 92 post-prime)

**Supplementary Figure 2. Correlation betseen the VNT_90_ or SARS-CoV-2-S1-specific IgG titers and ACE2 binding inhibitions. (A)** Correlation between VNT_90_ (Log_2_) against Wuhan, Delta (B.1.617.2), and Omicron (BA.1), and ACE2 binding inhibition (%) of 1:100 diluted sera from all animals of G1 (white circle), G4 (black triangle), and G5 (grey square) at week 94 (**B)** Correlation between SARS-CoV-2-S1-specific IgG titer against Wuhan and Omicron (B.1.617.2), and ACE2 binding inhibition (%) of 1:100 diluted sera from all animals of G1 (white circle), G4 (black triangle), and G5 (grey square) at weeks 92 and 94. Correlation between SARS-CoV-2 S1WU-specific (white circle) (**C**) or S1OM-specific (grey circle) (**D**) IgG titer, and ACE2 binding inhibition (%) of 1:100 diluted sera among sera from all animals of B. The lines represent the regression line of all samples and each symbol represents an individual mouse. Correlation analysis and calculation of Spearman’s correlation coefficients was performed using GraphPad Prism v9.

