## Supplementary figures and images for "Second Boost of Omicron SARS-CoV-2 S1 Subunit Vaccine Induced Broad Humoral Immune Responses in Elderly Mice"

### Supplemental Fig 1 and 2

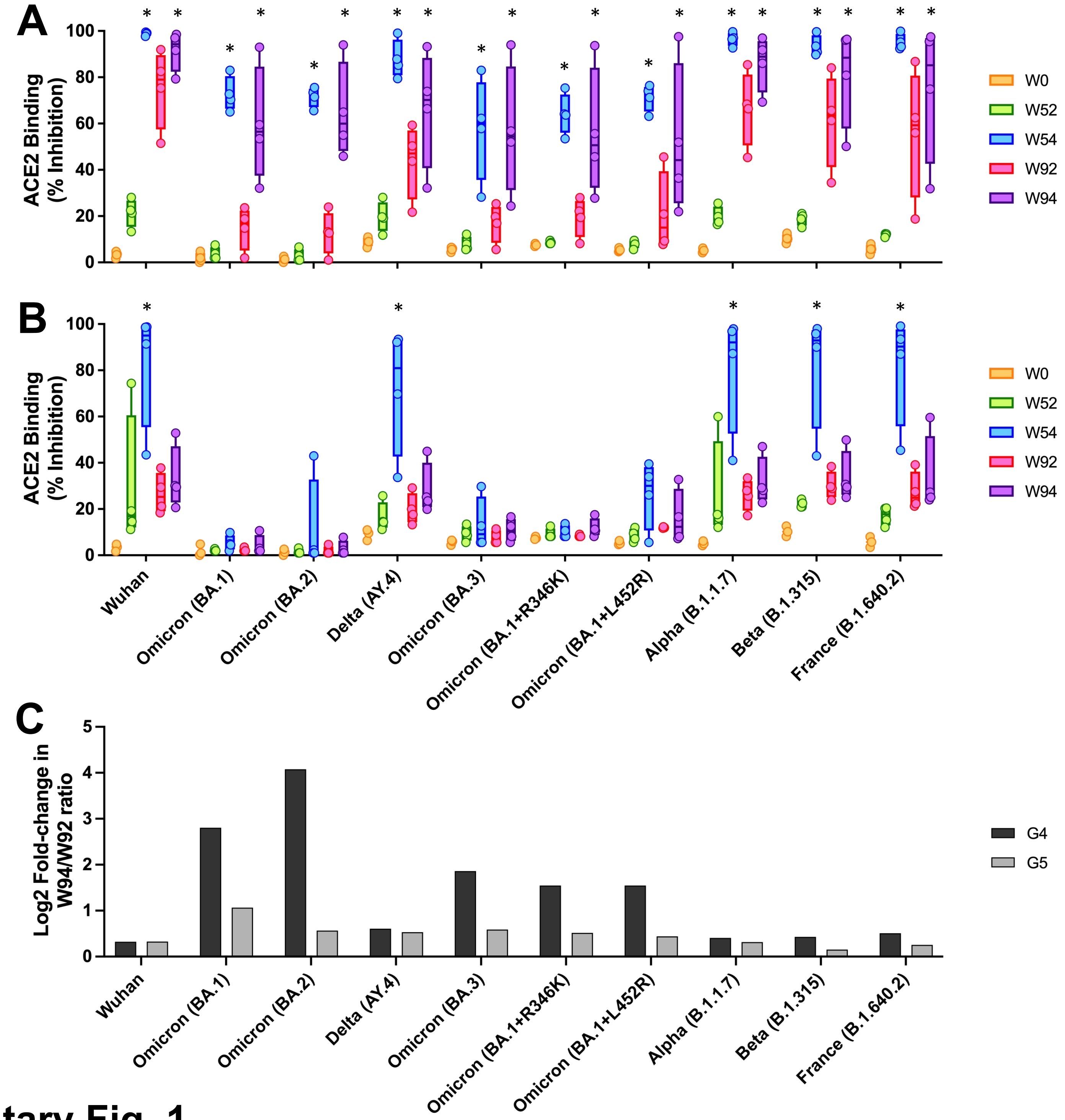


Supplementary Fig. 1

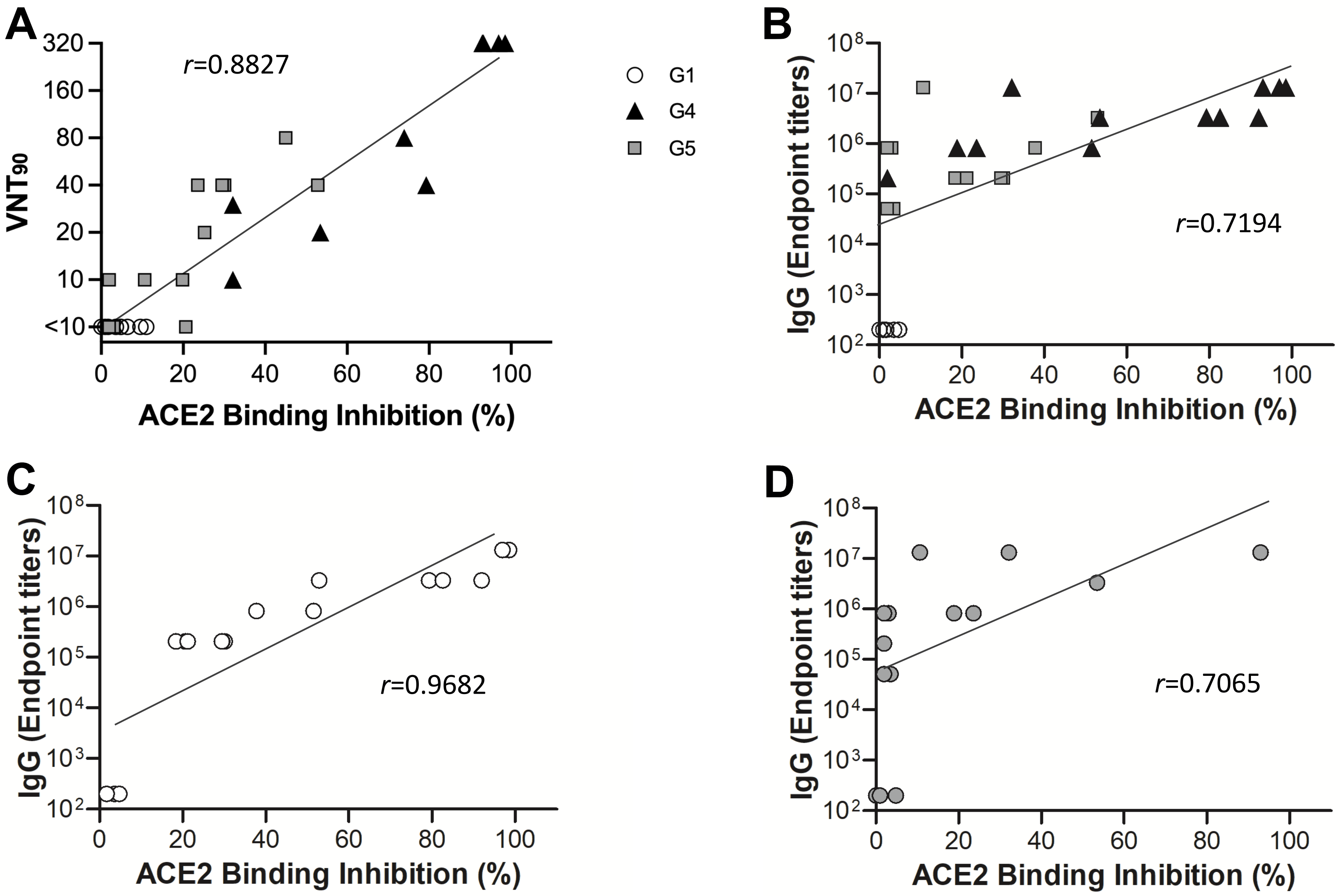


**Supplementary Fig. 2**
